# Exercise conditioned plasma dampens inflammation via clusterin and boosts memory

**DOI:** 10.1101/775288

**Authors:** Zurine De Miguel, Michael J. Betley, Drew Willoughby, Benoit Lehallier, Niclas Olsson, Liana Bonanno, Kaci J. Fairchild, Kévin Contrepois, Joshua E. Elias, Thomas A. Rando, Tony Wyss-Coray

**Affiliations:** Department of Neurology and Neurological Sciences, Stanford University School of Medicine, Stanford, CA 94305, USA; Neurosciences Graduate Training Program, Stanford University School of Medicine, Stanford, CA 94305, USA; Department of Chemical and Systems Biology, Stanford Medicine, Stanford, CA 94305, USA; The Veterans Affairs Palo Alto HealthCare System, Palo Alto, CA 94305, USA; Department of Genetics, Stanford University, 300 Pasteur Drive, Stanford, CA 94305, USA; Glenn Center for the Biology of Aging and Department of Neurology and Neurological Sciences, Stanford University School of Medicine, Stanford, CA 94305, USA; Wu Tsai Neurosciences Institute, Stanford University, Stanford, CA, USA

## Abstract

Physical exercise seems universally beneficial to human and animal health, slowing cognitive aging and neurodegeneration. Cognitive benefits are tied to increased plasticity and reduced inflammation within the hippocampus, yet little is known about the factors and mechanisms mediating these effects. We discovered “runner” plasma, collected from voluntarily running mice, infused into sedentary mice recapitulates the cellular and functional benefits of exercise on the brain. Importantly, runner plasma reduces baseline neuroinflammatory gene expression and prominently suppresses experimentally induced brain inflammation. Plasma proteomic analysis shows a striking increase in complement cascade inhibitors including clusterin, which is necessary for the anti-inflammatory effects of runner plasma. Cognitively impaired patients participating in structured exercise for 6 months showed higher plasma clusterin levels, which correlated positively with improvements in endurance and aerobic capacity. These findings demonstrate the existence of anti-inflammatory “exercise factors” that are transferrable, benefit the brain, and are present in humans engaging in exercise.

---

Physical activity evokes profound physiological responses in multiple tissues in species from fish, to birds, rats, mice and humans ^1–6^. It is widely accepted and promoted as a method of improving human health, including brain health ^7–9^. Exercise interventions in people of various ages with or without neurodegenerative diseases, or brain damage, have been shown to improve cognitive function ^9,10,11,12^. Neuroinflammation is a common feature of these conditions and potential mediator of the cognitive impairment associated with them ^13–16^. Studies in mouse models of aging and neurodegenerative diseases, such as AD and Parkinson disease have linked long-term voluntary wheel running with improved learning and memory, and decreased neuroinflammation ^17–20^. However, how exercise exerts these beneficial effects on the brain is poorly understood. It is possible that the physical exertion of muscle or lung during exercise may result in the secretion of factors from these or other tissues which subsequently signal to the brain to reduce neuroinflammation. Indeed, physical exercise increases levels of dozens of proteins in plasma, many of which are likely released from muscle tissue and thus named myokines ^2,21^. Some of these myokines, such as insulin-like growth factor-1 (IGF-1) ^22^, vascular endothelial growth factor (VEGF) ^23^ and platelet factor 4 (PF4) ^24^ increase following exercise and have been shown to increase hippocampal neurogenesis (Extended Table 1). However, it is unknown whether exercise conditioned plasma contains the factors benefitting the brain, whether these factors are directly transferrable through plasma, how such factors impact the brain, and what the key factors are.

## Runner plasma infusions induce neuroplasticity, improve learning and memory and regulate transcription of genes associated with the acute inflammatory response in the hippocampus

Given the well-known effects of voluntary wheel running in activating hippocampal neuroplasticity and neurogenesis in rodents ^25,26^, we tested the efficacy of plasma from exercising mice (“runners”) to increase neurogenesis. For all studies hereafter, we used plasma pools obtained from 3-month-old, male, C57Bl/6 mice housed in pairs for 28 days with access to a running wheel, producing “runner plasma” (RP) or access to a locked wheel, producing “control plasma” (CP). We chose this particular age and timepoint as we observed the most significant increase in overall hippocampal granule cell proliferation (EdU^+^ cells) as well as recently-born doublecortin positive (DCX^+^) neuroblasts across different ages from 3–15 months (Extended Data Fig. 1a, b). We further observed that 28 days of running was sufficient to increase overall cell survival (BrdU^+^ cells), including neurons (NeuN^+^/BrdU^+^ cells), the number of sex determining region Y-box 2 (SOX2) positive and glial fibrillary acidic protein (GFAP) negative neural stem and progenitor cells (NSPCs) and astrocytes (GFAP^+^/BrdU^+^ cells) (Extended Data Fig. 1c). In order to test whether physiological levels of circulating factors in runner plasma are sufficient to exert the well-known neurogenic effects of exercise, we then collected, pooled, and dialyzed plasma from each group of mice and injected it retro-orbitally into age-matched non-runners (Fig. 1a). Importantly, repeated control injections of *saline* via retro-orbital vein under anesthesia did not significantly alter hippocampal cell proliferation and survival (Extended Data Fig. 2). Twenty-four hours after the final plasma injection, we examined hippocampal cell survival and proliferation of new granule cells in the dentate gyrus of CP and RP recipient mice by quantifying the number of surviving (BrdU^+^) or dividing cells (EdU^+^).Remarkably, recipient mice receiving RP showed an average 31% increase in total proliferating cells, 40% increase of DCX^+^ neuroblasts, and 28% increase of surviving cells compared with CP recipient mice (Fig. 1b, c). Notably, these results are very similar to the direct effects of running described above (Extended Data Fig. 1). Interestingly, we replicated these findings using RP from 28-day-runners, and compared it with the effect of RP injections from 7- and 14-day-runners. While RP from 7- and 14-day-runners significantly increased neuroblast counts, it did not increase cell proliferation and survival (Extended Data Fig. 3). In addition, RP induced a 2-fold expansion of surviving NSPCs compared with CP and promoted the maturation of such cells into surviving NeuN^+^ cells (Extended Data Fig. 1c). However, RP did not significantly increase the number of surviving mature neurons and, instead, lead to a 2-fold increase in the survival of newborn astrocytes (Fig. 1c).

**Figure 1.**
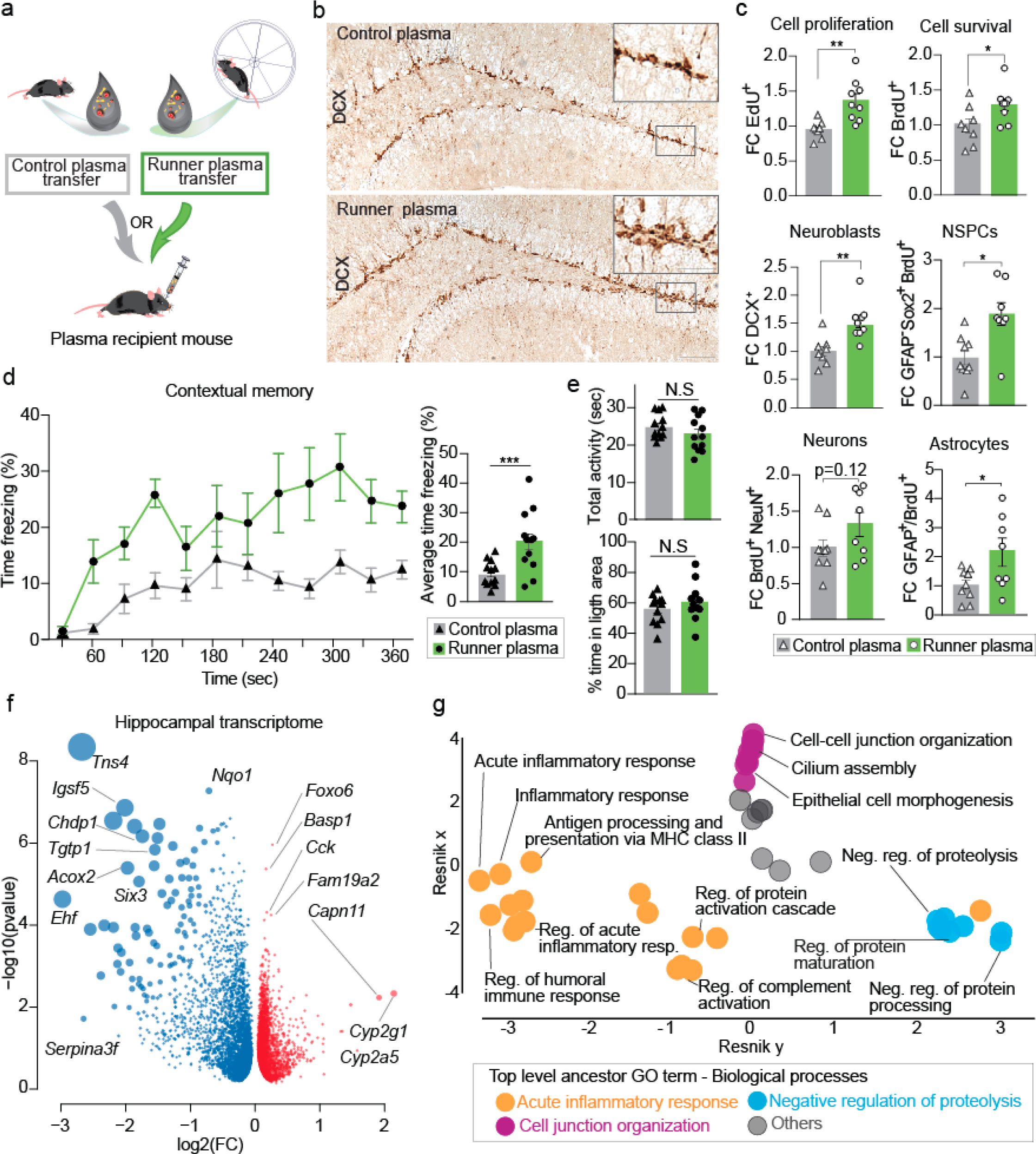
Runner plasma infusions induce neuroplasticity, improves learning and memory and regulates transcription of genes associated with the acute inflammatory response in the hippocampus. **a**, Plasma from running male mice (3-4 months of age.) was collected and transferred to matched aged and sex non-running mice, once every 3 days for 28 days. BrdU was administered 3 days before plasma administration and EdU 24 hours before sacrifice. Hippocampus was dissected and processed for transcriptome profiling using RNA sequencing and immunohistochemical detection of antigens. **b**, Representative images of immunolabeled DCX_+_ cells in the dentate gyrus of mice treated with control or runner plasma for 28 days. Bottom image scale bar 100 µm, insert 50 µm. **c**, Fold change per dentate gyrus (DG) of fluorescent immunolabeled EdU_+_ cells, BrdU_+_ cells, DCX_+_ cells, NeuN_+_/BrdU_+_ cells, GFAP_−_/Sox2_+_/BrdU_+_ cells and GFAP_+_/BrdU_+_ cells. (n=8-9 per group) **d**, Percentage of freezing behavior in response to the context associated with the fear stimulus in CP and RP recipient mice. (n=12 per group) **e**, CP and RP infused mice show comparable total activity and percentage of time spent in the dark arena. (n=12 per group) **f**, Volcano plot showing differentially expressed genes in the hippocampus of mice injected with CP or RP. Downregulated genes (blue); upregulated genes (red). Size of dot is the product of log_2_FC and −log_10_ p-value. (n=8 per group) **g**, Hierarchical networks of the abundance of gene ontology (GO) terms related to biological processes. GO terms correspond to the differentially expressed genes with treatment of CP versus RP, as shown in (f). Means ± s.e.m; unpaired Student’s two-tailed *t* test; N.S. not significant, * *P* < 0.05, ** *P* < 0.01 and *** *P* < 0.001

**Figure 2.**
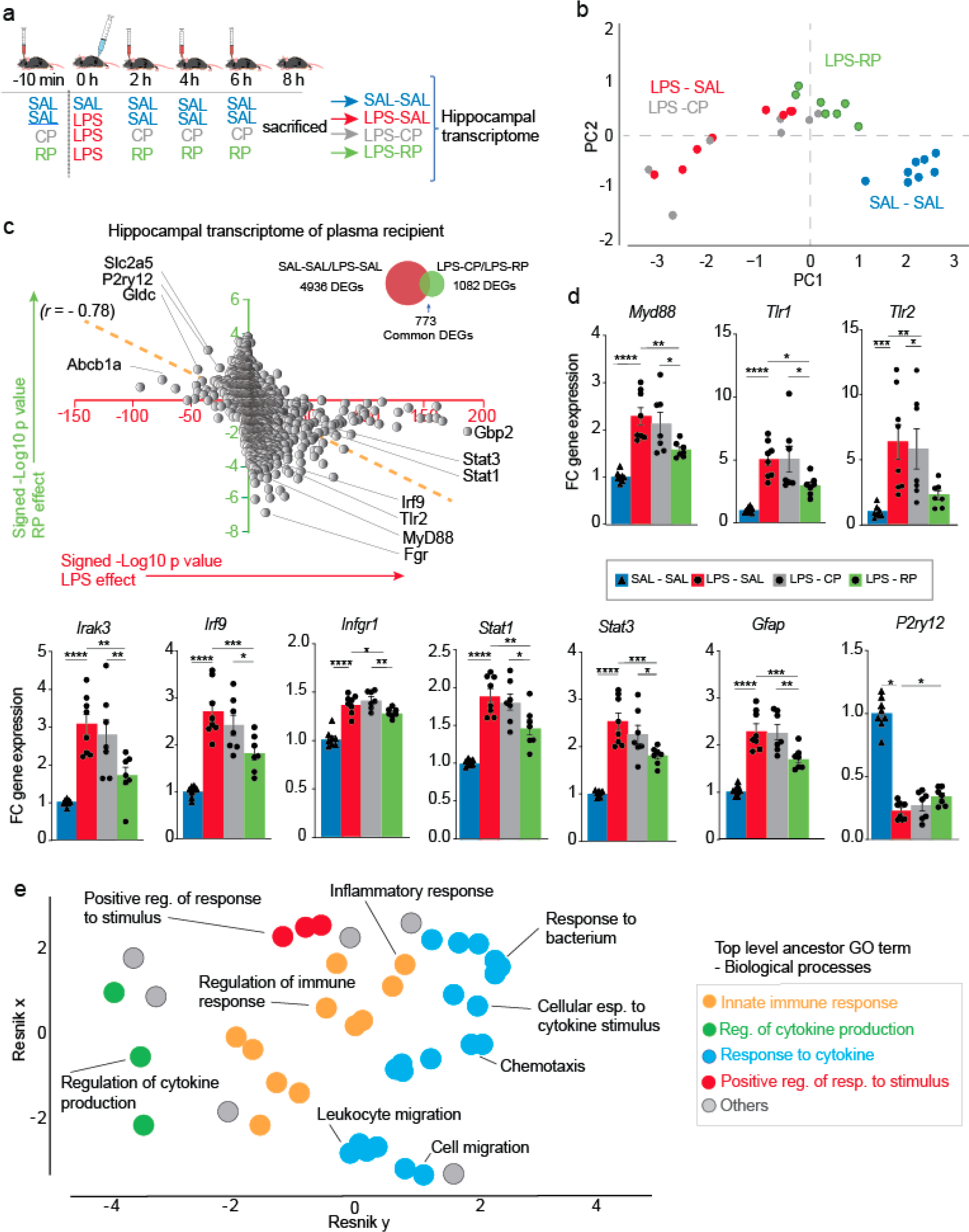
Runner plasma infusions reverse the neuroinflammatory effects of LPS on the hippocampus. **a**, Male mice (3-4 months of age) injected with LPS were treated with saline, runner plasma or control plasma (SAL – LPS, LPS – CP or LPS - RP). An additional control group received saline for all injections (SAL – SAL). Hippocampus was dissected and processed for transcriptome profiling using RNA sequencing. (n=7-8 per group). **b**, PCA analysis of common DEGs induced by LPS (SAL-SAL (blue) vs. SAL-LPS (red) and by RP treatment (LPS-CP (grey) vs. LPS-RP (green)). (n=7-8 per group). **c**, Correlation of signed −log_10_ p value changes of DEGs with LPS (SAL-SAL vs. SAL-LPS) and those induced with RP treatment (LPS-CP vs. LPS-RP). Venn diagram shows the proportion of DEGs induced by LPS in the hippocampus in relation to the DEGs induced by treatment with RP. (n=7-8 per group). **d**, Graphs show fold changes of relative gene expression of indicated genes as measured by PCR. (n=7-8 per group) **e**, Hierarchical networks of the abundance of gene ontology (GO) terms related to biological processes. GO terms correspond to DEGs in the hippocampus of LPS injected mice and treated with CP or RP. Means ± s.e.m; One-way ANOVA and Bonferroni post-hoc; * *P* < 0.05, ** *P* < 0.01, *** *P* < 0.001 and **** *P* < 0.0001

**Figure 3.**
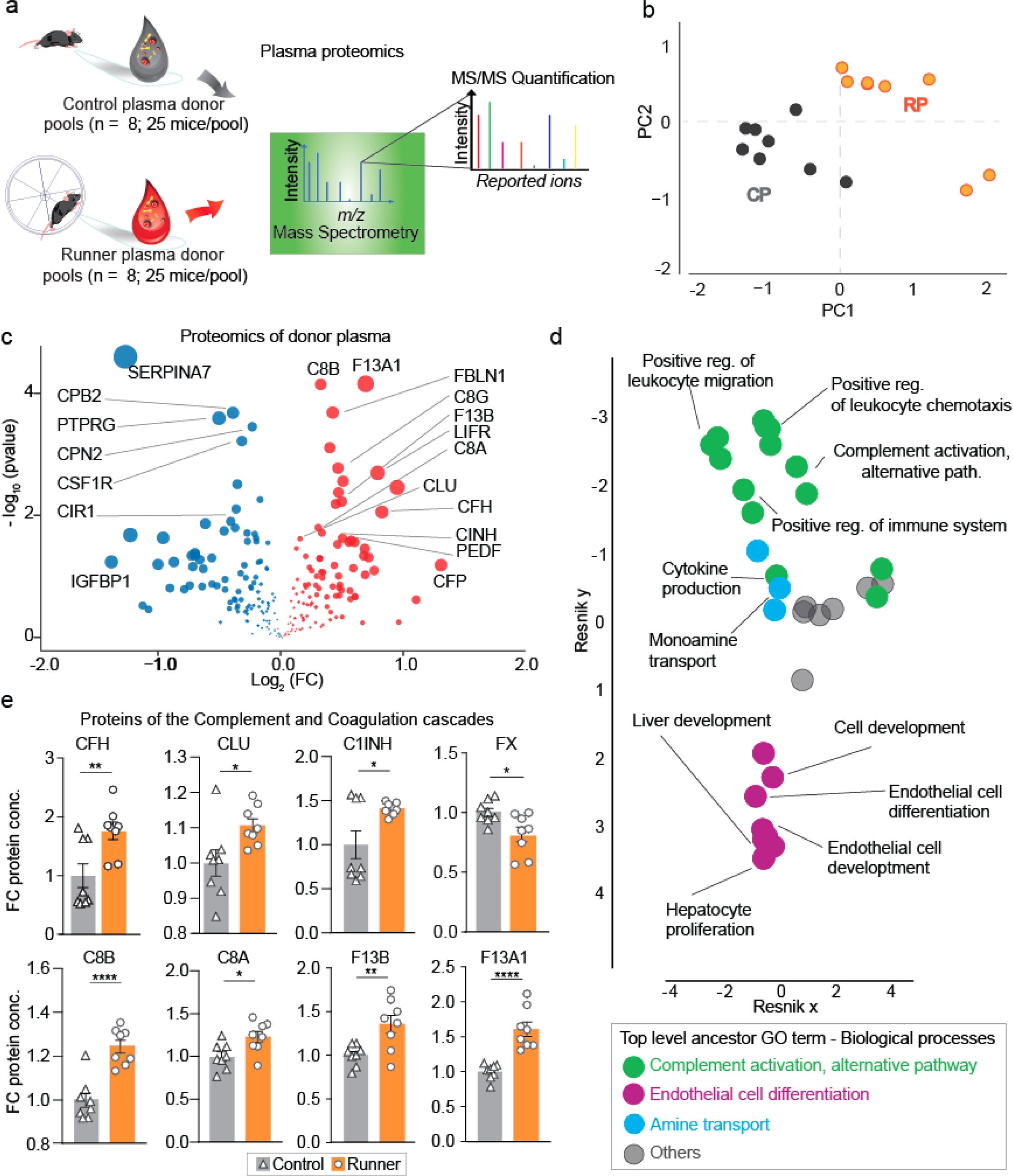
Running significantly alters levels of complement and coagulation pathway proteins in plasma. **a**, Eight pools of runner plasma (n = 25 male mice 3-4 months of age per pool) and control plasma (n = 25 male mice 3-4 months of age per pool) were collected, dialyzed and analyzed by TMT-LC-MS/MS. **b**, Classification by PCA analysis using plasma proteins detected by TMT-LC-MS/MS and significantly changed with running (p<0.05). Data was Log_2_ transformed and missing values were imputed by using the mean of each group. Control plasma (grey); runner plasma (orange). (n=8 per group) **c**, Graph showing proteins significantly changed (*P* < 0.05) in RP or CP. Downregulated proteins (blue); upregulated proteins (red). Size of dot is the product of log_2_FC and −log_10_ p-value. (n=8 per group) **d**, Hierarchical networks of the abundance of gene ontology (GO) terms related to biological processes of the proteins changed with 28 days of running. **e**, Representative plasma proteins of the complement and coagulation pathways significantly changed with running. (n=8 per group) Means ± s.e.m; unpaired Student’s two-tailed *t* test; * *P* < 0.05, ** *P* < 0.01, *** *P* < 0.001 and **** *P* < 0.0001

Remarkably, injection of RP, but not CP, led to a prominent increase in contextual learning and memory in a fear conditioning paradigm (Fig. 1d), consistent with studies reporting that exercise can improve cognitive function ^8,9^ and that increased hippocampal neurogenesis is linked to improved learning and memory ^27,28^. In contrast, we observed no difference in freezing behavior between CP and RP infused mice in response to auditory and visual cues in this behavioral paradigm (Extended Data Fig. 4). Importantly, mice receiving RP or CP showed comparable activity and anxiety levels in a light/dark arena (Fig. 1e).

**Figure 4.**
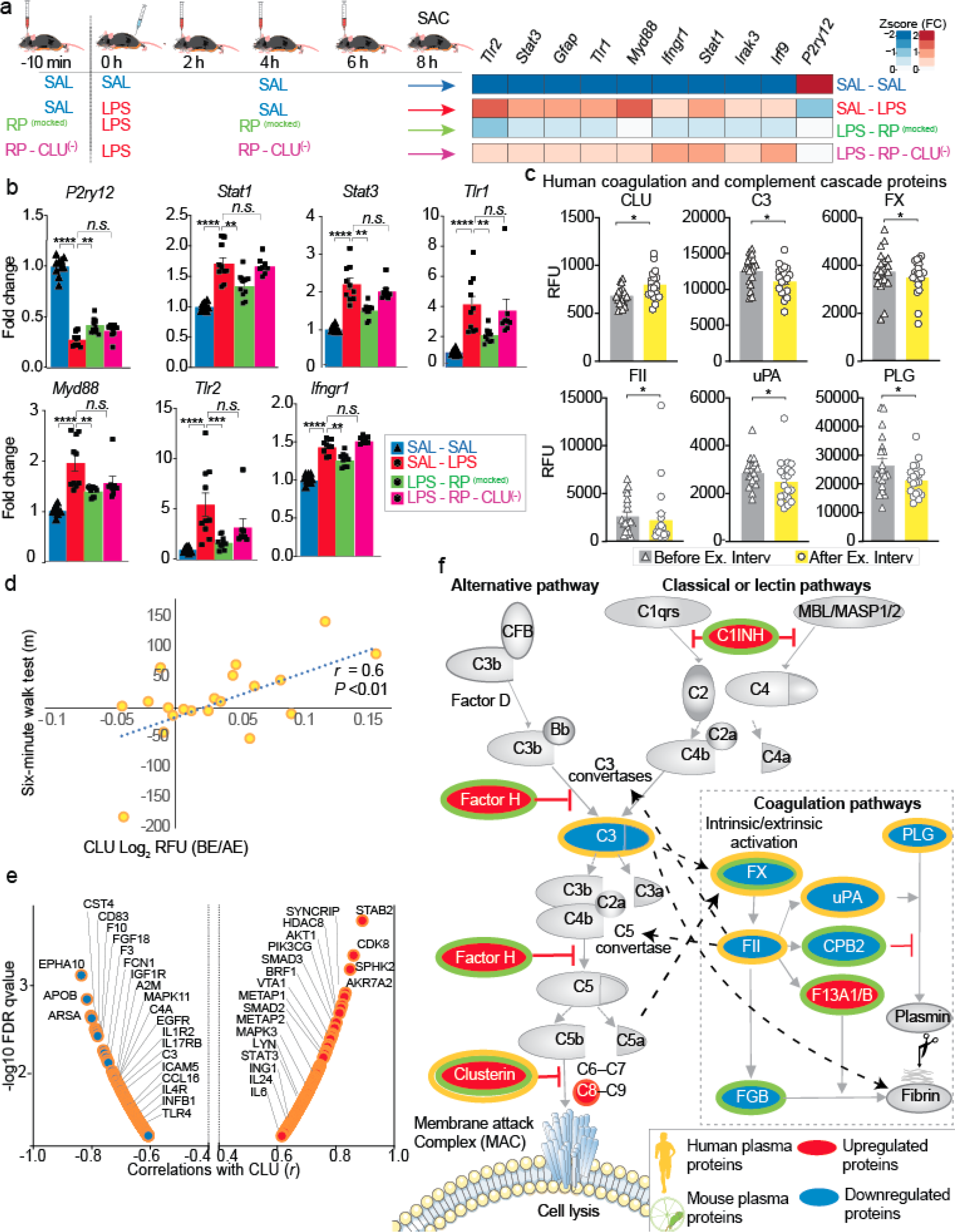
Runner plasma mediates the anti-inflammatory response to LPS in the hippocampus via clusterin. **a**, Male mice (3-4 months of age) were injected with LPS and treated with saline (LPS – SAL), runner plasma (LPS – RP) or runner plasma without CLU (LPS – RP – CLU_(−)_). An additional control group received saline for all injections (SAL – SAL). Heat map of selected inflammatory gene markers in the hippocampus of all the experimental groups. (n=8-10). **b**, Plots show individual values for each group on indicated genes measured by qPCR. (n=8-10). One-way ANOVA and Bonferroni post-hoc; n.s. not significant, * *P* < 0.05, ** *P* < 0.01, *** *P* < 0.001, **** *P* < 0.0001. **c**, Plasma CLU in humans before and after 6 months of exercise intervention measured in relative fluorescent units (RFU) as determined by SOMA-scan measurements (n=20 patients). Paired Student’s two-tailed *t* test; * *P* < 0.05 **d**, Plot shows correlation of changes in plasma CLU (before exercise (BE)/ after exercise (EA) with number of meters walked during the six-minute walk test (after exercise – before exercise). (n=20 patients) **e**, Graph shows correlations and the −log_10_ FDR q-value of changes in plasma CLU (before exercise (BE)-after exercise (EA)/(EA) with immune related plasma proteins ((BE-EA)/EA) (n=20 patients). **f**, Schematic representation of the significantly changed (*P* < 0.05) plasma proteins of complement and coagulation cascades in humans and mice after exercise. Dotted black arrows indicate relationship between factors of the complement and the coagulation system.

To understand the overarching effects of plasma factors on the hippocampus at the molecular level we analyzed the transcriptome of mice treated with CP or RP using RNA sequencing (RNAseq). We observed 1974 significant differentially expressed genes (DEGs), of which 61% were downregulated and 39% upregulated with RP treatment (Fig. 1f and Supplementary table 1). Genes implicated in cell migration (*Tns4)* ^29^, adhesion (*Igsf5)* ^30^ and epithelial cell differentiation and proliferation (*Ehf*) were downregulated while genes implicated in hippocampal synaptic function during learning and memory formation (*Foxo6*) ^31^ and immune system and plasticity (*Fam19a2*) ^32^ were upregulated. Intriguingly, analysis of functional categories of the top 250 DEGs defined by Gene Ontology (GO) and summarized into non-redundant hierarchical terms by ReviGO, pointed to a down regulation of the *acute inflammatory response* as the most significantly affected biological pathway, followed by upregulation of *cell junction organization* and *negative regulation of proteolysis* (Fig. 1g). These findings indicate that factors in plasma from running mice are largely sufficient to reproduce the effects of running on hippocampal neurogenesis, to improve learning and memory and to downregulate the baseline expression of inflammatory genes in the hippocampus.

## Runner plasma infusions reverse the neuroinflammatory effects of LPS on the hippocampus

Peripheral lipopolysaccharide (LPS) administration, the major component of the outer membrane of Gram-negative bacteria, is commonly used to model the neuroinflammatory conditions associated with neurodegenerative diseases by inducing central neuroinflammation without damaging the brain parenchyma ^33,34^. Because running plasma infusion downregulates immune and inflammatory genes (Fig. 1f, g) we tested if RP infusions could reduce acutely induced neuroinflammation in mice systemically inoculated with LPS (Fig. 2a). Administration of LPS resulted in 4,902 DEGS (Supplementary table 2) in the hippocampus. Strikingly, treatment with RP, but not CP, reversed the expression of a large number of these genes (*r* = −0.78) such that the hippocampal transcriptome of RP treated mice had greater similarity to that of mice that did not receive LPS (Fig 2 b-c). For example, RP reduced the expression of genes, which were validated by qPCR (Fig 2 d, Supplementary table 3), of the toll like receptor (TLR) signaling pathway (*Myd88*, *Tlr1* and *Tlr2* and *Irak3*), the interferon pathway (*Irf9*, Infgr*1*, Stat*1* and *Stat3)*, as well as the glial fibrillary acidic protein (*Gfap*), a sensitive indicator of astrocyte activation. Pathway analysis of the most significant 500 DEGs pointed to *response to cytokine and regulation of the innate immune response* and *regulation of the response to stimulus and cytokine production* (Fig 2e). Together, these results demonstrate that RP effectively reverses the neuroinflammatory response to LPS in the hippocampus.

## Running significantly alters levels of complement and coagulation pathway proteins in plasma

While the preceding studies analyzed the effects of RP on the brains of recipient mice we wanted to identify the key factors mediating these effects. We thus made large pools of plasma from 28-day-runners and controls (Fig. 3a) and subjected them to unbiased detection of proteins via tandem mass tag (TMT) isobaric labeling of proteins and synchronous precursor selection-based MS3 (SPS-MS3) mass spectrometry. We identified 235 unique proteins simultaneously detected across the two conditions; of these, 49 proteins were significantly changed with running (23 downregulated and 26 upregulated) and PCA analysis segregated the two groups according to their treatment (Fig. 3b, c; Supplementary table 4). We validated these results using a shotgun proteomic approach (Extended Data Fig. 5). Biological pathway analysis pointed to *activation of the alternative pathway of the complement system and endothelial cell differentiation* (Fig 3d). Remarkably, proteins of the complement and coagulation pathways represent 26% of the differentially expressed proteins. Indeed, it has become clear over the past few years that these two pathways are functionally related ^35,36^. While complement is generally activated by infection and coagulation is activated by tissue damage, factors of each pathway triggers the activation of the other. For example, the complement factors C3a and C5a can directly activate platelets and induce tissue factor (TF) expression, respectively, via interaction with their cognate receptors ^37,38^, while coagulation factors such as thrombin can activate C5 directly^39^ and C3 indirectly ^40^. Our proteomic analysis pointed to upregulation of three inhibitors of the complement system, factor H (FH), complement 1 inhibitor (C1INH) and clusterin (CLU), possibly indicating a downregulation of complement activation (Fig. 3e). Additionally, we observed a concerted increase in C8 alpha and gamma chains of the C8 component of the terminal pathway for the formation of membrane attack complex (MAC) (Fig. 3e), possibly acting as a MAC formation inhibitor by preventing C5b7 from attaching to the target cell membrane ^41^. Within the coagulation cascade, we observed a downregulation of factor X (FX/10), fibrinogen (FGB) and carboxypeptidase B2 (CPB2) and an upregulation of factors F13A1 and F13B, which promote fibrin clot formation and stabilization by crosslinking fibronectin to the fibrin clot. FX is the first component of the thrombin pathway, which ultimately leads to the conversion of FGB into a fibrin clot. In the same vein, CPB2, also downregulated, is known as a thrombin-activatable fibrinolysis inhibitor, inhibiting plasmin mediated degradation of fibrin products and suggesting that the clotting cascade is overall inhibited in runners. On the other hand, the upregulation of F13 alpha and beta chains, which are known to stabilize fibrin, could indicate that compensatory mechanisms for normal homeostasis are active as well. Together, these findings demonstrate a profound effect of running on the complement and coagulation cascades in the blood.

## Clusterin is a key protein for the anti-inflammatory effects of RP on the hippocampus

To determine if complement inhibitory proteins were critical for the anti-inflammatory effects of RP on the hippocampus we immunodepleted clusterin and factor H from pools of RP. We selected clusterin because of its potent inhibitory effects on terminal pathway activation ^42^ (Fig. 3c, e) and Factor H because of its inhibitory effects of the alternative pathway accelerating the decay of the C3 convertase (C3b,Bb) and inactivation of C3b ^43^. For comparison, we also depleted glycoprotein pigment epithelium-derived factor (PEDF) and leukemia inhibitory factor receptor (LIFR), which are known to have anti-inflammatory functions ^44,45^ and were among the top regulated proteins with exercise (Fig. 3b). Mice were injected systemically with LPS and treated with intact RP (mock immunodepleted with IgG control antibodies) or RP lacking clusterin, Factor H, PEDF or LIFR and hippocampi of mice were analyzed 8h later for expression of immune and inflammatory genes by qPCR. Remarkably, while control depleted RP reduced hippocampal inflammation, depletion of the complement inhibitor clusterin in RP largely abrogated the anti-inflammatory properties of RP. Depletion of Factor H had a lesser effect on RP’s anti-inflammatory capacity and depletion of PEDF or LIFR had no effect, indicating that plasma clusterin is a key factor in reducing brain inflammation via exercise-conditioned plasma (Fig. 4a, b. Extended Data Fig. 6). To determine if clusterin and complement/coagulation cascades were changed in response to exercise in humans, we analyzed the plasma proteome, using a commercial aptamer-based proteomic platform, of twenty Veterans with amnestic Mild Cognitive Impairment (aMCI) (Extended Data Table 2) before and after a 6-month exercise intervention. Here, we report not only a significant increase in plasma clusterin levels in response to exercise but a positive correlation between changes in clusterin levels and changes in aerobic capacity and endurance (Fig. 4c, d). We also observed significant reductions in complement factor 3 (C3), FX, coagulation factor II (FII/Prothrombin), uPA, and plasminogen (PLG) in response to exercise in these individuals (Fig. 4c). Furthermore, changes in clusterin plasma levels induced by exercise significantly correlated with changes in 260 out of 1304 detected plasma proteins (Fig. 4e; Supplementary table 5). For example, clusterin correlates positively with CDK8 and MAPK3, both of which have been implicated in transcriptional activation of cell cycle progression ^46,47^. Clusterin was also positively correlated with IL-24 and IL-6 which can act as anti-inflammatory ^48–50^. Interestingly, we observed that clusterin correlated negatively with the complement factors C3, C4a and the coagulation factor FX as well as with proteins of the toll like receptors and interferon families (Fig. 4e). Enrichment analysis using Gene Ontology of biological processes revealed that proteins correlated with clusterin participate in the *regulation of the immune response.* In line with the effects observed in mice (Fig. 4f), these results suggest a concerted downregulation of the complement and coagulation pathways which correlates with functional improvements in mice and humans.

## Discussion

Here, we show that changes in plasma as a result of physical exercise, and the magnitude of these changes, are sufficient to recapitulate the beneficial effects of exercise on neurogenesis and memory and to reduce neuroinflammation in the hippocampus of non-exercised mice. We identify clusterin as a novel exercise-induced plasma protein mediating anti-inflammatory effects of exercise plasma on the hippocampus and show that clusterin levels correlate positively with functional benefits of exercise in humans. Our unbiased transcriptomic analysis of brains from mice receiving runner plasma indicates that runner plasma regulates immune and inflammatory pathways in the hippocampus. Furthermore, runner plasma also has profound anti-inflammatory effects in a model of acute neuroinflammation.

Based on multiple experiments and the use of independent proteomic methods in human and mice, our findings indicate that running induces concerted changes in plasma proteins within the complement and coagulation systems. While the anti-inflammatory effects of exercise has been recognized, it has typically been linked to changes in the secretion of immune modulatory cytokines from muscle and fat tissue, the mobilization of regulatory T cells from lymphoid organs, or the reduction of inflammatory monocytes in the blood ^51^.To our knowledge, changes in the complement and coagulation cascades have not been implicated in the effects of exercise in reducing neuroinflammation within the brain. Importantly, we identified clusterin as a novel exercise-induced plasma protein mediating the anti-inflammatory effects of exercise plasma on the hippocampus. Furthermore, we show that clusterin levels correlate positively with functional benefits of exercise and a reduction of inflammatory plasma proteins in humans.

Clusterin, a complement inhibitor, is a highly conserved and ubiquitous glycoprotein, with high expression levels in multiple immune cells as well as in hepatocytes (Extended Data Fig.7). Hepatocytes are a major source of proteins in the complement and coagulation cascades ^52^ making it tempting to speculate that exercise prompts the liver to induce the observed changes reported here. In addition to its role in inhibiting inflammation via the complement system^53^ or the NF-kappaB pathway ^54^, polymorphisms in the *CLU* gene are linked to AD ^55,56^. Current evidence suggest that during neurodegeneration, there is an upregulation of complement activation, and attenuating its activation by inhibiting C1q or C3 results in lower neuroinflammation, Aβ plaques and synapse loss ^57^ ^58^. In patients with AD and MCI, plasma clusterin levels are increased ^59^ possibly as a compensatory mechanism ^60^. It is possible that clusterin’s peripheral inhibition of the complement system induced by exercise might contribute to the beneficial effects of exercise in neurodegenerative diseases or aging, by similarly inhibiting the complement system in the brain. How clusterin exerts anti-inflammatory effects on the brain needs further investigation.

We also demonstrate that runner plasma treatment improves cognitive function and recapitulates the effects of exercise on neural stem cell activity. Recent studies have found that neuroinflammation negatively affects neurogenesis and cognition ^61,62^. Given that plasma complement inhibitor CLU reduces neuroinflammation, we speculate that this pathway could mediate the improvement in memory and the increase in neural stem cell activity induced by runner plasma treatment. Collectively, the findings presented here provide new insight into the mechanisms of how exercise benefits the brain. Our results offer new avenues to develop therapies based on proteins induced by exercise that are capable of reducing neuroinflammation and improve cognitive function.

## Acknowledgements

We thank members of the Wyss-Coray lab for their support, D. Berdnik for excellent technical assistance, H. du Bois and S. Shuken for sharing their protocols, A. Yang and J. Marschallinger for useful comments on the manuscript, and H. Zhang and K. Dickey for excellent laboratory management. This work was funded by the US National Institute on Aging (AG047820; T.W.-C. and T.A.R.), the US Department of Veterans Affairs (Research Career Scientist Award IO1 BX001319; T.W.-C.), the Department of Defense (W81XWH-12-1-0584; K.J.F.), the Alzheimer’s Association (NIRG-15-362171; J.K.F.) and by the Marie Curie Foundation n-273487 (GCs-CNS-IS) awarded to ZDM. The contents supported by this funding do not represent the views of the VA or the United States Government.

## Author Contributions

T.W.-C., T.A.R, Z.D.M, and M.B. designed and conceived the experiments; Z.D.M. and M.B. developed an initial paradigm of plasma transfer from runner to non-runner mice and studied the effect of plasma on neural stem cell activity. Z.D.M. and D.W. performed experiments to generate plasma pools, carried out animal treatments and to process brain tissue and plasma samples for molecular and protein analyses. D.W. performed and analyzed behavioral experiments under the supervision of Z.D.M. L.B. and Z.D.M performed and analyzed sequencing experiments. B.L. executed statistical analyses and visualization of protein and gene data sets. K.J.F. performed experiments with humans and collected plasma samples, N.O, J.E.E. and K.C. carried out Mass Spectrometry analyses. Z.D.M., D.W. and T.W.-C. wrote the manuscript with input from T.A.R.. T.W.-C. and Z.D.M. supervised the study.

## Methods

### Animals

C57Bl/6J male mice (#000664, Jackson Laboratory) were purchased from Jackson Labs and housed in groups of 2-4 mice prior to any exercise or plasma injections. C57Bl/6J male mice used in aging studies were obtained from the National Institute on Aging (NIA). All mice were housed at the Palo Alto VA animal facility under a 12hr:12hr light:dark cycle with dark hours between 6:30PM – 6:30AM. All experiments were performed in accordance with institutional guidelines approved by the VA Palo Alto Committee on Animal Research.

### Voluntary Exercise

All exercise experiments were performed using activity cages with a running wheel installed. Activity cages with a computer-monitored running wheel were purchased from Lafayette Instrument Co. (Lafayette, IN). After arrival, mice were housed in groups under standard laboratory conditions for 7 days prior to the start of exercise. Subsequently, running mice were housed in the activity cages in pairs with free access to unlocked wheels and control mice were housed in pairs in standard animal facility cages.

### Processing and administration of plasma

Three to four-month-old C57Bl/6 male mice were housed in pairs either in activity cages with access to a running wheel or in standard housing cages (controls) for 28 days. Running and control mice were sacrificed between 6:30-9:00 am. Mice were anesthetized with 18 μL of a 2.5% solution of Avertin per gram body weight (2,2,2-tribromoethanol: Sigma Aldrich Cat# T48402, 2-methyl-2-butanol: Sigma Aldrich Cat# 2404860) in preparation for perfusion or plasma collection. Blood was collected from the right heart ventricle with 30uL of 250mM EDTA (ThermoFisher, Cat# 15575020) and centrifuged at 4°C for 15 min at 1000 × g to collect plasma. Plasma from 20-25 exercised or control mice was pooled together and dialyzed using cassettes (Slide-A-Lyzer Dialysis Cassettes, 3.5K MWCO, 3-12 mL) and then frozen at −80°C. For plasma transfer experiments, non-running mice were injected retro-orbitally with 200 μL of plasma per injection.

### BrdU, EdU, and LPS preparation and administration

Both 5-bromo-2’-deoxyuridine (BrdU) and 5-ethynyl-2-deoxyuridine (EdU) were resuspended in sterile PBS and injected intraperitoneally at 50mg/kg. For long-term cell survival assays, BrdU was administered 72, 48 and 24 hours before the start of exercise or plasma treatment. To assay proliferation, EdU was administered 12 hours before sacrifice. Lipopolysaccharide (LPS, Sigma, Cat# L4391) was resuspended in sterile saline at 10μg/mL and injected at 100μg/kg. Control mice were treated with an equivalent volume of saline. Sample size was estimated using historical data to provide sufficient power (80%) to assess effects of RP on neurogenesis (n = 8).

### Tissue processing and Immunohistochemistry

Mice were anesthetized as previously described and transcardially perfused with 20 mL of 1M phosphate buffered saline (PBS). Right hemispheric hippocampal tissue was snap frozen in liquid nitrogen and stored at −80 °C for future transcriptomic or proteomic analysis. Left hemispheres were stored in 4% paraformaldehyde at 4 °C for 48 hours then switched to 30% sucrose solution for an additional 48 hours in preparation for brain sectioning. Serial coronal sections of the hippocampus (40 µm thick) were collected using a freezing microtome (Leica SM2010R) and stored in cryprotective medium (40% PBS, 30% glycerol, 30% ethylene glycol) at −20 °C until immunostained. Immunohistochemistry was performed using standard techniques described previously ^63^ using free-floating sections. For primary antibody information see Extended data table 3. All sections stained for BrdU or EdU were pre-treated with 3M HCL for 15 minutes at 37 °C before incubation with primary antibody. The Click-iT Plus EdU AlexaFluor® 555 Imaging Kit (ThermoFisher, Cat# C10638) was used for EdU detection. For total DCX immunoreactivity cell counts, a biotinylated anti-goat secondary antibody (rabbit, 1:1000, Vector) was used, and cells were then visualized using a Vectastain® Elite® ABC kit (Vector, Cat# PK-6100) and diaminobenzidine (Sigma-Aldrich, DAB, Cat# D5905). All fluorescent secondary antibodies were diluted at a concentration of 1:200 and incubated with sections for 2 hours in TBS-T (0.01 M Tris HCl pH 7.4 + 0.15 M NaCL + 5 × 10^4 mL/L Tween 20) at room temperature. All fluorescent secondary antibodies were Alexafluor® 488, 555, or 647 antibodies (ThermoFisher) raised in donkey against the appropriate target animal. Nuclei were fluorescently labeled with Hoechst 33342 (1:2000, Sigma).

### Imaging and cell quantification

Images of immunofluorescent sections were obtained, quantified, and analyzed by a blinded-observer to the treatment. Fluorescent microscopy was performed using a Zeiss 710 Confocal Laser Scanning Microscope (Carl Zeiss MicroImaging, Zeiss EC Plan-Neofluar, 20×, 40×, and 63×/0.05) with the detection pinhole set at 1 Airy Unit. Optical sections were taken at 2 μm intervals, and the obtained z-stack images were processed and analyzed with the National Institutes of Health ImageJ software. The number of labelled cells were manually counted in six representative sections. One hundred or more BrdU or EdU immunoreactive cells were evaluated per mouse and the total number of cells was corrected for number of optical slices or percent area.

### Behavior

Behavioral testing began the day following final plasma injections. Experimenters were blinded to the treatment. *SmartCage* (AfaSci, SmartCage system): Animals were allowed to explore the cage freely for 30 minutes, location, total locomotion and time spend in light/dark areas was evaluated using the CageScore software (AfaSci). *Fear Conditioning:* Fear conditioning, contextual testing, and cued testing took place in sound-attenuating cages equipped with steel shocking floor ((CleverSys Inc., Cat# CSI-BOX-STD and CleverSys Inc., Cat# CSI-CHM-FLR-M). Behavior in cages was recorded by low-light camera and stimulus, shock, and freezing behavior analysis was automated using the FreezeScan system and software (CleverSys Inc.). During fear conditioning training, mice were habituated to shock cages for 2 min and then were subjected to two pairings of a 30 sec, 2 kHz auditory cue (70 dB, cage enclosed) and a white light cue with a co-terminating 2 sec foot shock (0.6 mA). The intratrial interval was 2 min, and 2 min after the final shock mice were returned to their home cage. The following day mice underwent contextual and cued recall testing, with an intertrial interval of 2 hours. Contextual testing was performed in the same cages used during the training trial. Animal freezing behavior was tracked over a period of 6 min in the absence of any auditory or visual cues. Cued testing was performed in a different context, using cages with altered odor (3% acetic acid), floor texture, and wall shape. Mice were subjected to the same pattern of auditory and visual cues as during training (without a shock pairing) and were returned to home cages 2 min after final cues. Sample size was estimated using the literature and historical data to provide sufficient power (80%) to assess effects of RP on behavior (n = 12).

### RNA preparation and microfluidic quantitative real time PCR

RNA was isolated from hippocampal tissue using RNeasy mini kits (Qiagen, Cat# 74104) according to manufacturer’s instruction. A primer pair panel containing 10 neuroinflammatory related genes + 10 housekeeping genes for microfluidic quantitative real time PCR (qRT-PCR) was designed as previously described ^64^. Creation and pre-amplification of cDNA was performed using 100ng RNA and reverse transcription and pre-amplification kits from Fluidigm (Cat#100-6301) following the manufacturer’s protocol. Sample and assay mixes were created using cDNA diluted 1:5 and appropriate primer pairs according to manufacturer instructions. The 96.96 Dynamic Array IFC for Gene Expression chip (Fluidigm, Cat# BMK-M-96.96) was then loaded and mixed using a Biomark IFC Controller HX (Fluidigm, Cat# BMK-IFC-HX) before processing and data acquisition using a Biomark HD Real-Time PCR System (Fluidigm, Cat# BMKHD-BMKHD). Transcript fold change was calculated using ΔΔCt values based on experimental controls.

### RNA-sequencing

RNA was isolated from hippocampal tissue as previously described and quality was validated using an Agilent 2100 Bioanalyzer (Stanford PAN Facility). RNA was converted into cDNA using a SMART-seq® v4 Ultra Low Input Kit for Sequencing (Takara Bio, Cat# 634894) according to the manufacturer’s protocol. cDNA was then fragmented, normalized, and pooled using a Nextera XT library preparation kit (Illumina, Cat# FC-131-1096) according to manufacturer’s instructions. Sequencing was performed on Illumina Novaseq 6000 to obtain pared-end 150bp reads. Reads were aligned to the mouse genome using the STAR GeneCounts function and differential expression analysis was performed using the R DESeq2 package ^65^ using built-in algorithms.

### Plasma Immunodepletion Studies

Antibodies to target proteins were coupled to superparamagnetic beads using a Dynabeads® Antibody Coupling Kit (ThermoFisher, Cat# 14311D) according to manufacturer’s instructions. Antibodies were conjugated to beads at a ratio of 10μg antibody/mg beads. Antibody-conjugated beads were stored at 50mg/mL in sterile PBS at 4 °C until use. To deplete plasma, 32μL of antibody-conjugated bead suspension was added to 800μL dialyzed plasma (2mg beads/mL plasma) and incubated for 1 hr at room temperature and 23 hours at 4 °C. Depleted plasma was separated from beads using a magnetic rack and stored at −80 °C until use. For antibody detailed information see Extended data table 3.

### Plasma Albumin and IgG depletion

In all plasma MS experiments, albumin and immunoglobulin were depleted from 15 μL of dialyzed plasma using an immunoaffinity column (PROTIA Sigma ProteoPrep Immunoaffinity Albumin & IgG Depletion Kit, Cat# PROTIA-1KT). Depleted plasma and albumin/IgG fractions bound to the immunoaffinity columns were eluted separately and stored at −20°C in Axygen Maxymum Recovery microcentrifuge tubes (Corning/Fisher, Cat# MCT-175-L-C) until further processing.

### Mass Spectrometry (MS)

#### Cleanup, reduction, and alkylation

A total of 100μg protein was diluted in phosphate buffered saline (pH 7.4, ThermoFisher, Cat# 10010) to a final volume of 300 μL. Diluted samples were denatured at 95°C for 10 min, cleaned with 5 μL Benzonase (Millipore, Cat# 70664) for 30 min at 37°C, reduced for 30 min at 45°C with 15 μL of 200 mM dithiothreitol (DTT, ThermoFisher, Cat# R0861), and incubated at room temperature for 30 min with 15 μL of 400 mM iodoacetamide (Sigma, Cat# I1149). Alkylation reaction was quenched using 30 uL of 200 mM DTT.

#### Single-Pot Solid-Phase-enhanced Sample Preparation (SP3)

The SP3 method used in this study was adapted from previous methods ^66^. Equal parts hydrophobic and hydrophilic Sera-Mag carboxyl magnetic beads (GE Healthcare, Cat# GE65152105050250 and GE45152105050250) was added to each sample at a ratio of 15 μg beads per 1 μg protein. Samples were acidified with 10 μL of 1% formic acid (FA, Fisher/Pierce, PI28905) and 100% LC/MS grade acetonitrile (ACN, Fisher/Pierce, Cat# PI51101) was added to reach at least 50% of the total volume. After mixing and incubating at room temperature for 8 min, protein-laden beads were aggregated and collected using a magnetic rack (ThermoFisher, Cat# 12321D). Samples were then washed twice with 200 μL of 70% ethanol (Sigma, Cat# E7023) and once with 180 μL of ACN,. Finally, proteins were incubated overnight (~16 hrs) at 37°C in 50 μL of a 0.1μg/μL solution of Trypsin/LysC (Fisher/Promega, V5073) in 50 mM HEPES buffer (ThermoFisher, Cat# 15630). Following digestion, samples were sonicated (Fisher, Cat# 15-337-22) for 10 min.

Samples for subsequent tandem-mass-tag (TMT) processing (see below) were then magnetically separated from beads and stored at −80°C until TMT labeling. ACN was added to samples to be used for shotgun MS to reach at least 95% of the total volume. After incubation for 8 min at room temperature, supernatant was discarded and beads were washed with 180 μL ACN. Beads were reconstituted in 50 μL of 2% dimethyl sulfoxide (DMSO, ThermoFisher, Cat# 85190) and sonicated for 5 min, supernatant containing peptides was retained. 5% of FA was added to the samples and immediately processed through a STAGE tip cleanup.

#### Tandem-mass-tag (TMT) labeling and Fractionation

TMT labeling was achieved using a TMT10plex Isobaric Label Reagent Set (ThermoFisher, Cat# 90111) according to the kit’s directions. A global standard was created to enable a comparison between 10plex sets by taking aliquots of equal volume from each sample and combining them. Portions of each sample were used to check ion distributions and labeling efficiency. The rest was combined in adjusted ratios with the other labeled samples in their respective TMT10plex set, lyophilized in a centrifugal vacuum concentrator (Labconco, Cat# 7810010), and resuspended in 5% FA for STAGE tip cleanup. Some TMT labeled peptide samples were separated into 8 fractions using high pH reversed-phase fractionation columns (ThermoFisher Scientific, Cat# 84868). Lyophilized peptide samples were resuspended in 0.1% trifluoroacetic acid (TFA) and fractionated according to manufacturer’s instructions. Samples were then lyophilized and resuspended in 0.1% FA for LC-MS/MS analysis.

#### STAGE tip cleanup and peptide recovery

STAGE tips were made by packing unfiltered 200μL micropipette tips with C18 Empore extraction disks (VWR, Cat# 55004-098). For each equilibration and wash step, 120 μL of solvent was added to the STAGE tip and centrifuged at approximately 2500 × g for 2 min such that solvent levels were just above the filter layers, preventing filters from drying out. Each STAGE tip was incubated for 5 min with methanol (Sigma, Cat# 34885), then washed twice with 50% ACN + 5% FA and twice with 5% FA. Peptide samples were added to equilibrated STAGE tips and centrifuged into filters as previously described. STAGE tips containing peptides were washed twice with 5% FA, and peptides were then eluted in 100μL of 50% ACN + 5% FA, lyophilized, and stored at −80°C until resuspension in 0.1% FA for LC-MS/MS analysis. Retention time calibration peptides (ThermoFisher Scientific, Cat# 88321) were spiked into resuspended shotgun samples at 125 fmol/μL.

#### Mass spectrometry, data acquisition, and analysis

Peptides were analyzed on either an Orbitrap Fusion or an Orbitrap Fusion Lumos Mass Spectrometer (ThermoFisher). Peptides were separated by capillary nanoflow reversed phase chromatography on a reversed-phase column (100 μm inner diameter, packed in-house with ReproSil-Pur C18-AQ 3.0 m resin (Dr. Maisch)) using either two or three hour long LC-gradients. The LC buffers used were mobile phase A: 0.1 % formic acid in water and mobile phase B: 0.1% formic acid in acetonitrile. Full MS scans were acquired in the Orbitrap mass analyzer at a resolution of 120,000 (FWHM) in a data-dependent mode. The AGC targets were 4*10^5^ and the maximum injection time for FTMS1 were 50 ms. *Shotgun label free MS:* The most intense ions were then selected for sequencing and fragmented in the Orbitrap mass analyzer using higher-energy collisional dissociation (HCD) with a normalized collision energy of 30% and resolution of 15,000 (FWHM). Monoisotopic precursor selection was enabled and singly charged ion species and ions with no unassigned charge states were excluded from MS2 analysis. Ions within ±10 ppm m/z window around ions selected for MS2 were excluded from further selection for fragmentation for 30 sec. AGC target were 5*10^4^ and maximum injection time of 250 ms. Shotgun label free MS2 spectra were analyzed using MaxQuant v1.6.3.4 and searched against the Uniprot Mouse sequence database. MaxQuant defaults were used for all unmentioned settings. Carbamidomethylation of cysteine was set as a static modification and protein N-acetylation and methionine oxidation were variable modifications. A false discovery rate of 1% at the peptide and protein levels was set, and only peptides with at least 7 residues were included. At least one unique or razor peptide in the protein group was required for protein identification. Whenever possible, the matching between runs feature in MaxQuant was used to identify peptides present in MS1 spectra but not chosen for MS2 fragmentation and sequencing in a given sample based on matching retention times and mass with other samples in which that peptide was identified. MaxQuant’s label free quantification (LFQ) algorithm was used to quantify peptides and proteins in shotgun samples with the fast LFQ mode enabled. *TMT MS:* the most intense ions were selected from full MS scans in top speed mode for sequencing using collision-induced dissociation (CID) and the fragments were analyzed in the ion trap. The normalized collision energy for CID was 35% at 0.25 activation Q. For MS2 the AGC targets were 2*10^4^ and the maximum injection time set to 50 ms for the Fusion while for the Fusion Lumos the AGC targets were 1*10^4^ and the maximum injection time 30 ms. Monoisotopic precursor selection and include charge state 2-5 (Fusion) or 2-7 (Fusion Lumos) were enabled. Singly charged ion species and ions with no unassigned charge states were excluded from MS2 analysis. Ions within ±10 ppm m/z window around ions selected for MS2 were excluded from further selection for fragmentation for either 45 s (Fusion) or 90 s (Fusion Lumos). Following each MS2 analysis, the most intense fragment ions were selected simultaneously for HCD MS3 analysis. For MS3 analysis acquired on the Fusion an isolation window of 2 at a resolution of 60,000, with m/z scan range 100-500, normalized collision energy of 65%, AGC target 5*10^4^ and maximum injection time of 118 ms were used. For MS3 analysis on the Fusion Lumos an isolation window of 1.2 m/z, normalized collision energy of 65%, resolution of 60,000, m/z scan range 120-500, AGC target of 1*10^5^ and maximum injection time of 90 ms were used. The raw data for TMT labeled samples was analyzed using Proteome Discoverer v2.2.0.388 (ThermoScientific). Precursor mass tolerance was ±10 ppm and fragment mass tolerance was ±0.6 Da. Static modifications were carbamidomethylation of cysteine (+57.021 Da) and TMT-labeled N-terminus and lysine (+229.163 Da). Differential modifications were oxidation of methionine (+15.995 Da), phosphorylation of serine, tyrosine and threonine (+79.9663), acetylation of protein N-terminal (+42.011 Da). Proteome Discoverer searched the spectra against the Uniprot Mouse database, including the identification of common contaminants using the SEQUEST algorithm. Percolator was applied to filter out false MS2 assignments at a false discovery rate of 1% at peptide and protein levels. For quantification, a mass tolerance of ± 20 ppm window was applied to the integration of reporter ions using the most confident centroid method. The average abundance of the top 3 peptides mapped to each protein was used to estimate protein abundances.

### Gene Ontology (GO) and REViGO Analyses

For RNA-seq and MS proteomic GO analysis, genes and proteins were ranked by their differential expression and LFQ abundance p values, respectively. For RNA-seq analysis, GO terms were annotated from the top genes and the GO terms enrichment was tested using the topGO R package ^67^, with remaining detected genes were kept in the analysis as background. GO term enrichment analysis for MS proteomics was performed analogously on the top proteins detected. The most significant GO terms were summarized into non-redundant hierarchical terms by their semantic similarity via ReviGO ^68^, using Resnik as the clustering algorithm and 0.7 for allowed similarity measure.

### Human exercise regimen

Twenty Veterans with amnestic Mild Cognitive Impairment (aMCI) participated in the experiment. Diagnosis was based on the National Institute on Aging/ Alzheimer’s Association criteria ^69^. Human samples were obtained prior to and after completion of a combined aerobic and resistance exercise training program. Veterans participated in thrice weekly exercise sessions for six months. All exercise sessions included the 5 minutes warm-up and cool-down periods prior to and following 30 minutes of continuous aerobic exercise (e.g., treadmill walking, stationary cycling, cycle ergometry) and 20 minutes of full-body resistance exercises. Target exercise intensities were 60-70% of maximum heart rate, as determined by cardiopulmonary treadmill testing. These studies were approved by Institutional Review Boards.

### Human plasma collection and proteome data acquisition

Human plasma was collected from the upper Arm by the Routine Blood Draws in the Clinical Pathology Laboratory at the Veterans Hospital of Palo Alto between July 2013 and 2017 and storate at −80°C. Samples were processed by SomaLogic Inc. (Boulder, CO) using the SOMAscan v3 assay ^70^. Samples were processed under Good Laboratory Practice ^71^ for intra-run normalization and calibration and passed all quality control requirements.

