## Extended data figure 1 for "Exercise conditioned plasma dampens inflammation via clusterin and boosts memory"

### Extended data 1

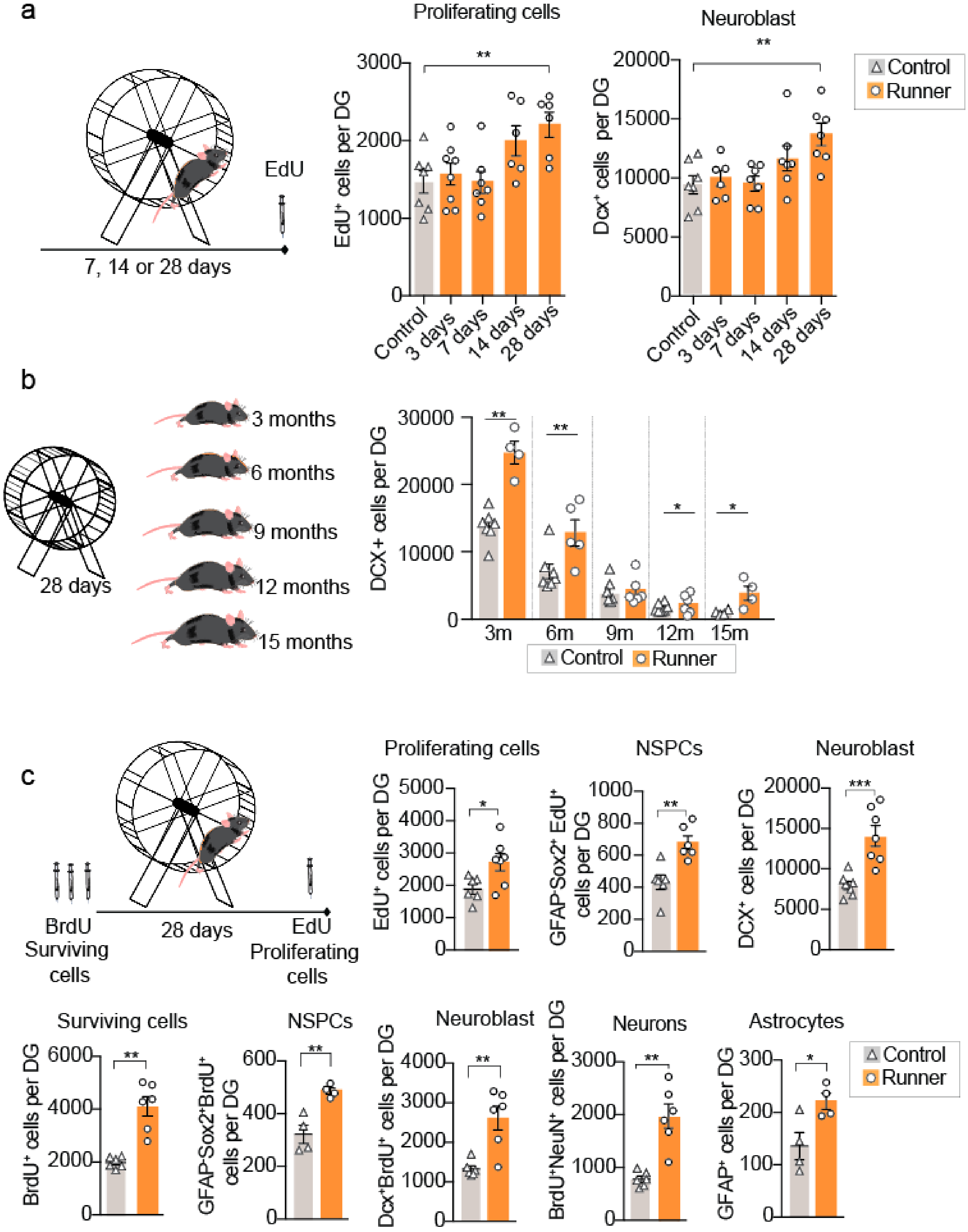

**Extended data 1. Twenty-eight days of voluntary wheel running increases hippocampal neural stem activity.**

**a**, Male mice at 3 months of age had access to a running wheel for 3, 7, 14 or 28 days while controls remain without access to running wheels. EdU was administered 24 hours before sacrifice. Plots show total number of cells per dentate gyrus (DG) of fluorescent immunolabeled $EdU^+$ cells and $DCX^+$ cells (n=6-7 per group).

**b,** Male mice at 3, 6, 9, 12 and 15 months of age had access to a running wheel for 28 days. Graphs show total number of $DCX^+$ cells per dentate gyrus. (n=4-8 per group)

**c,** Male mice at 3 months of age had access to a running wheel for 28 days. BrdU was administered 3 days before the exercise and EdU 24 hours before sacrifice. Graphs show total number of cells per dentate gyrus (DG) of fluorescent immunolabeled cells. (n=4-6)

Means ± s.e.m; unpaired Student's two-tailed *t* test; * $P < 0.05$, ** $P < 0.01$ and *** $P < 0.001$
