## Extended data figure 2 for "Exercise conditioned plasma dampens inflammation via clusterin and boosts memory"

### Extended data 2

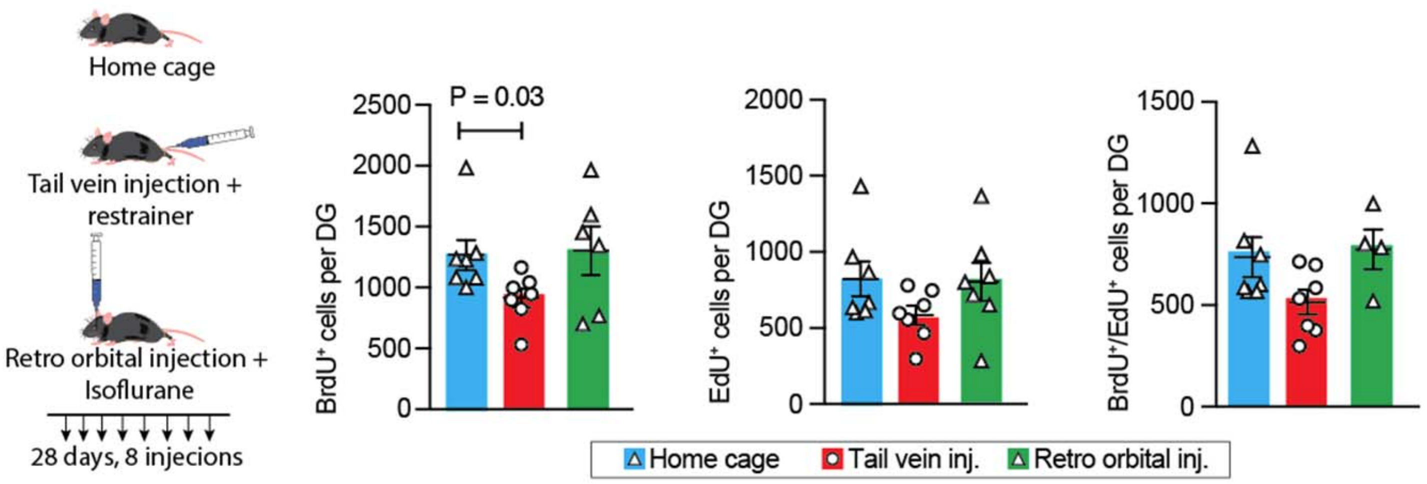


**Extended data 2. Control injections of saline via retro-orbital vein in combination with Isoflurane do not impair neural stem activity**

Mice at 3 months of age were injected with saline via the tail vein or the retro orbital vein with 200 µl of saline, every 3 days for 28 days. BrdU was administered 3 days before saline administration and EdU 24 hours before sacrifice and the hippocampus was dissected and processed for immunohistochemistry. Graphs show total number of cells per dentate gyrus (DG) of fluorescent immunolabeled cells.
