## Extended data figure 3 for "Exercise conditioned plasma dampens inflammation via clusterin and boosts memory"

### Extended data 3

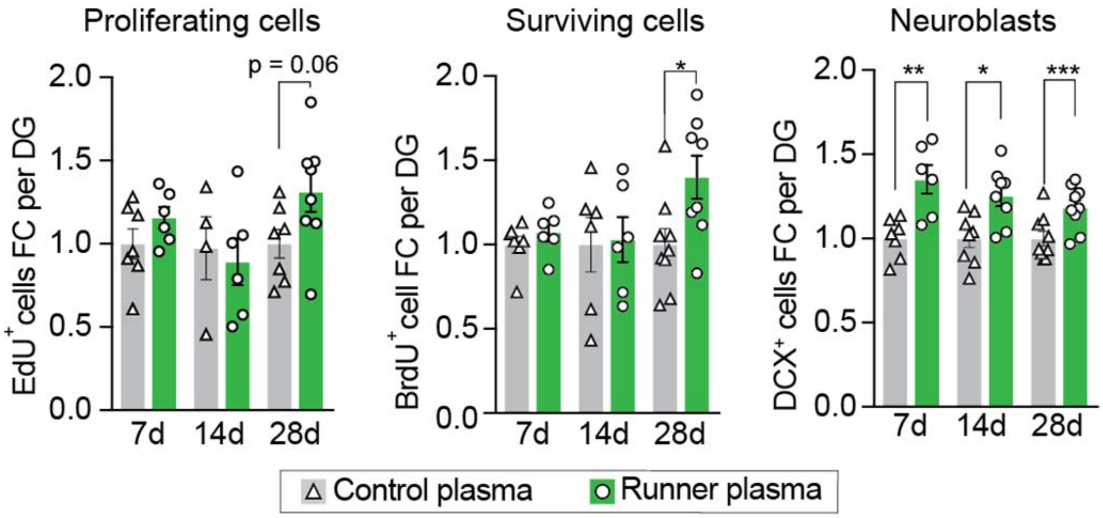


**Extended data 3**. **Runner plasma infusions from 28 days runners upregulate proliferation and survival of hippocampal new born cells.**

Plasma from running mice (3-4 months of age) that run for 7, 14 or 28 days was collected and transferred to matched aged non-running mice, once every 3 days 28 days. BrdU was administered 3 days before plasma administration and EdU 24 hours before sacrifice. Graphs show total number of cells per dentate gyrus (DG) of fluorescent immunolabeled $EdU^+$ cells, $BrdU^+$ cells and $DCX^+$ cells (n=6-9 per group).

Means ± s.e.m; unpaired Student's two-tailed *t* test; * $P < 0.05$, ** $P < 0.01$ and *** $P < 0.001$
