## Extended data figure 4 for "Exercise conditioned plasma dampens inflammation via clusterin and boosts memory"

### Extended data 4

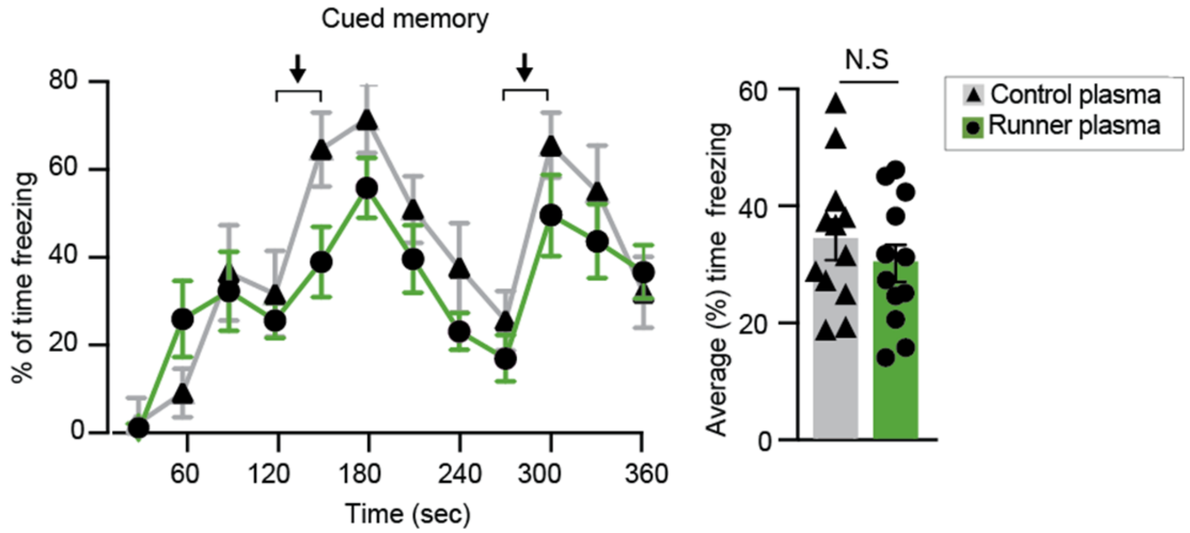


**Extended data 4. Runner plasma infusions do not significantly affect cued memory.**

Plasma from running mice (3-4 months of age) was collected and transferred to matched aged non-running mice, once every 3 days for 28 days. Mice were then tested for cued memory on the fear conditioning test. Graphs show percentage of freezing behavior in response to the light/tone cues associated with the fear stimulus in CP and RP recipient mice. (n=12 per group)

Means ± s.e.m; unpaired Student's two-tailed *t* test; N.S., no significant
