## Extended data figure 5 for "Exercise conditioned plasma dampens inflammation via clusterin and boosts memory"

### Extended data 5

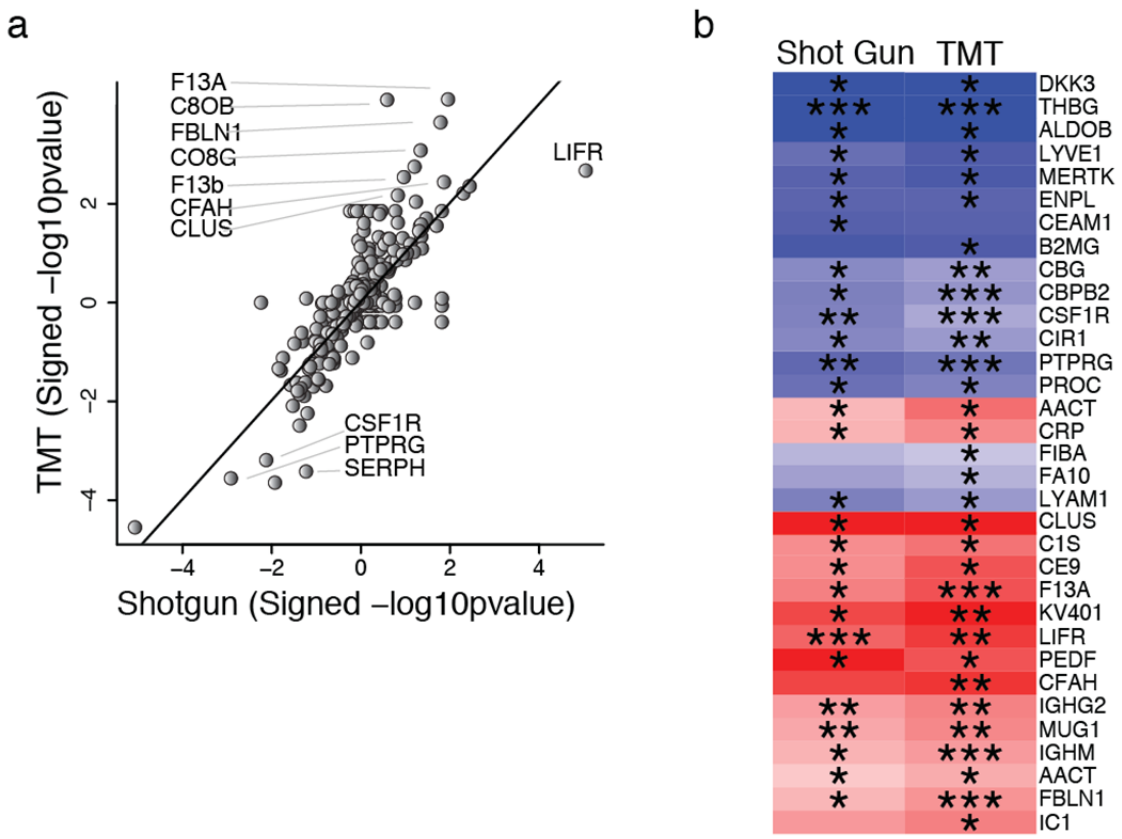


**Extended data 5. Validation of proteins captured with TMT-LC MS/MS using shotgun-LC MS/MS detection and analysis.**

**a**, Plot show the correlation of proteins significantly changed in plasma pools from runners and controls (n = 8 per group), in X-axis detected in TMT-LC MS/MS relative quantification, and in Y-axis detected in shotgun-LC MS/MS relative.

**b**, Heat map depicting the relative levels of the top differentially expressed plasma proteins detected with TMT- or shotgun- LC MS/MS.

Means ± s.e.m; Unpaired Student's two-tailed *t* test; * $P < 0.05$, ** $P < 0.01$ and *** $P < 0.001$
