## Extended data figure 6 for "Exercise conditioned plasma dampens inflammation via clusterin and boosts memory"

### Extended data 6

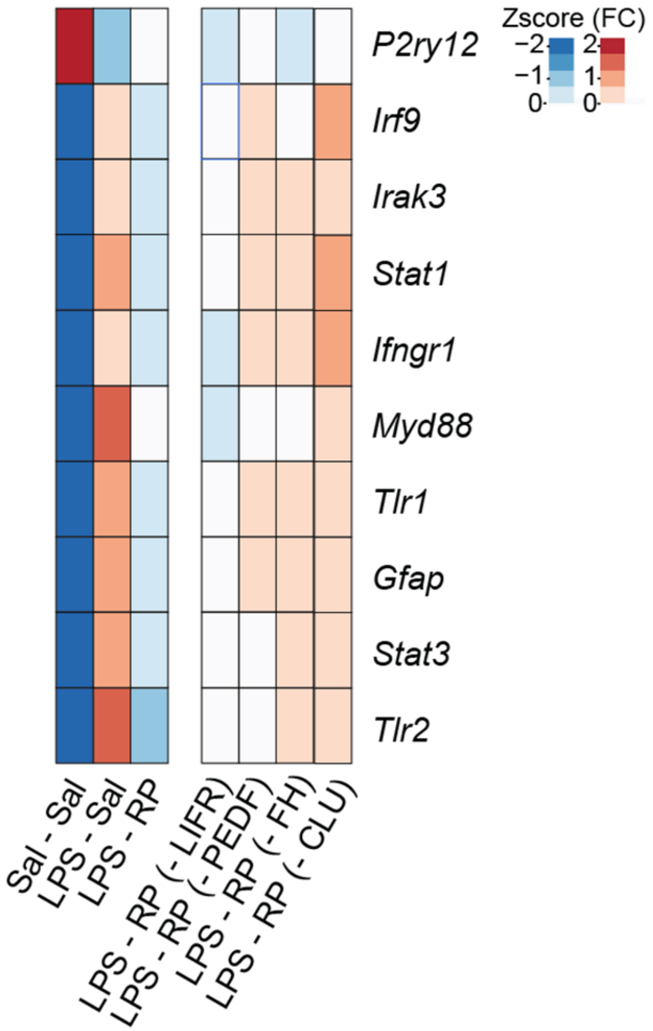


**Extended data 6. Immunodepletion of clusterin in runner plasma abrogates its anti-inflammatory properties on the hippocampus.**

Male mice (3-4 months of age) were injected with LPS and treated with saline (LPS – SAL), runner plasma (LPS – RP), runner plasma without CLU (LPS – RP – CLU), runner plasma without FH (LPS – RP – FH), runner plasma without LIFR (LPS – RP – LIFR)) or runner plasma without PDEF (LPS – RP – PDEF). Heat map of selected inflammatory gene markers in the hippocampus. Of all the immunodepleted groups, the LPS – RP – CLU group shows the highest Euclidean distance to the LPS – RP group.
