## Extended data figure 7 for "Exercise conditioned plasma dampens inflammation via clusterin and boosts memory"

### Extended data 7

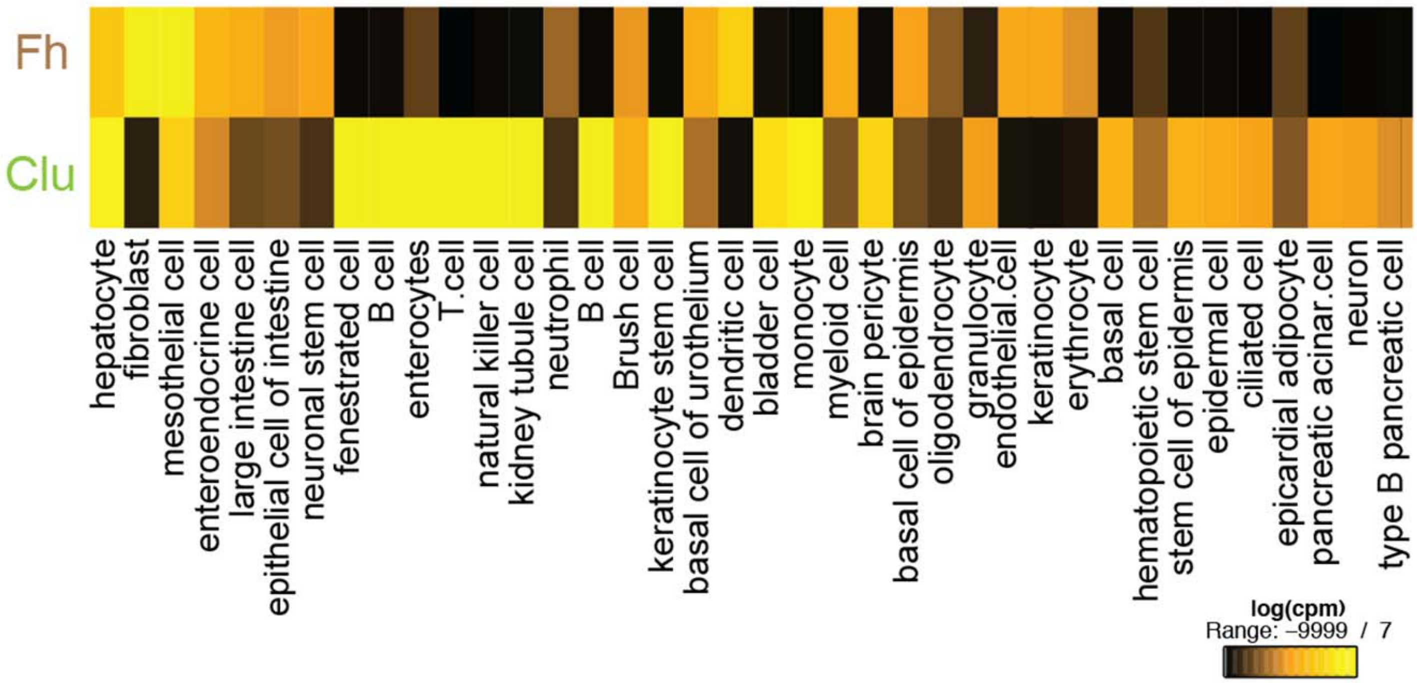


**Extended data 7. Expression of *Clu* and *Fh* in different cell types.**
Heat map showing expression of *Clu* and *Fh* in different cell types based on single cell gene expression in the Tabula Muris Atlas [1]
