## Extended data table 1 for "Exercise conditioned plasma dampens inflammation via clusterin and boosts memory"

| Name of molecule or cell | Change in the periphery with exercise | Is the molecule sufficient for the effect? | Is the molecule necessary for the effect? | Brain endpoint | Model system | Citation |
|---|---|---|---|---|---|---|
| **Insulin-like growth factor 1 (IGF-1)** | | YES<br>*In vivo;* IGF-1 subcutaneus infusion for 7 days increases **BrdU+ cells in the hippocampus**. | YES<br>*in vivo*; IGF-1 antiserum administered subcutaneously during 2 weeks of treadmill running reduces **BrdU+ cells in the hippocampus** | **Neuroplasticity** | Rat | Trejo et al. 2001 |
| | ↑ Increased in serum with long term training<br>↓ Decreased | | | | human | Koziris, L. P. et al. 1999<br>Eliakim, A. et al. 1996 and 1998 |
| **Vascular endothelial growth factor (VEGF)** | ↑ Wheel running | | YES<br>*in vivo*; overexpression of VEGF antagonist in the periphery during 7 days of voluntary wheel running decreased **BrdU+ cells** | **Neuroplasticity** | mouse | Fabel et al 2003<br>Laufs U et al. 2004. |
| **Cathepsin B (CTSB)** | ↑ CTSB plasma levels increase with 3, 14 or 30 day of running | | YES<br>*in vivo*: mice CTSB KO that run for 28 days showed no increase of **DCX+ cells in the hippocampus** | **Neuroplasticity** | mouse | Moon et al. 2016 |
| **Irisin (FNDC5)** | ↑ In plasma after 3 weeks of free wheel running | YES<br>*in vivo:* adenoviral overexpression of FNDC5 in the liver increases **BDNF in the hippocampus**. | | | mouse | Wrann et al. 2013<br>Bostrom P et al. 2012 |
| **PF4/platelets** | ↑ PF4 plasma levels increase in mice that ran for 1 day | | YES<br>*in vivo*; anti-platelet serum administered during 13 days of voluntary wheel running prevented the **increase of Ki67+ cells in the hippocampus** | **Neuroplasticity** | mouse | Leiter et al. 2019 |
| **β-endorphin** | | | YES<br>*in vivo*: β-E KO prevents the exercise-induced increase in **hippocampal Ki67+ cells** with 10 and 39 days of running,<br>NO<br>Doesn't affect exercise-induced increases in **DCX+ or BrdU+ cells.** | **Neuroplasticity** | mouse | Koehl et al 2008 |
| | ↑ Increase in long distance runners | | | | human | Colt et al. 1981 |
| **Lactate** | ↑ Increase with treadmill running | YES<br>Intraperitoneal lactate injections increased hippocampal lactate and **BDNF gene expression and protein** | YES<br>*in vivo*: intraperitoneal injections of lactate MCT1/2 inhibitor during exercise abolished exercise-induced **BDNF expression in the hippocampus** | **Neuroplasticity** | mouse | Hayek et al. 2019<br>Ferreira et al. 2007 |
| **Brain derived neurotrophic factor (BDNF)** | | YES<br>Intravenous BDNF after focal cerebral ischemia **reduces neurological deficit and infarct volume.** | | **Infarct volume** | rat | Schabitz et al 2000 |
| | ↑ Increase in plasma with treadmill running | | | | human | Cho et al. 2012 |

**Extended data table 1.**
Summary of the effects of *in vivo* peripheral interventions to manipulate proteins that have been proposed to be upregulated in the periphery with exercise and to have an effect in the brain[1-15].

### Extended data table 1 references


1 Bostrom, P. *et al.* A PGC1-alpha-dependent myokine that drives brown-fat-like development of white fat and thermogenesis. *Nature* **481**, 463-468, doi:10.1038/nature10777 (2012).

2 Cho, H. C. *et al.* The concentrations of serum, plasma and platelet BDNF are all increased by treadmill VO(2)max performance in healthy college men. *Neurosci Lett* **519**, 78-83, doi:10.1016/j.neulet.2012.05.025 (2012).

3 Colt, E. W., Wardlaw, S. L. & Frantz, A. G. The effect of running on plasma beta-endorphin. *Life Sci* **28**, 1637-1640, doi:10.1016/0024-3205(81)90319-2 (1981).

4 El Hayek, L. *et al.* Lactate Mediates the Effects of Exercise on Learning and Memory through SIRT1-Dependent Activation of Hippocampal Brain-Derived Neurotrophic Factor (BDNF). *J Neurosci* **39**, 2369-2382, doi:10.1523/JNEUROSCI.1661-18.2019 (2019).

5 Eliakim, A., Brasel, J. A., Mohan, S., Wong, W. L. & Cooper, D. M. Increased physical activity and the growth hormone-IGF-I axis in adolescent males. *Am J Physiol* **275**, R308-314, doi:10.1152/ajpregu.1998.275.1.R308 (1998).

6 Fabel, K. *et al.* VEGF is necessary for exercise-induced adult hippocampal neurogenesis. *Eur J Neurosci* **18**, 2803-2812 (2003).

7 Ferreira, J. C. *et al.* Maximal lactate steady state in running mice: effect of exercise training. *Clin Exp Pharmacol Physiol* **34**, 760-765, doi:10.1111/j.1440-1681.2007.04635.x (2007).

8 Koehl, M. *et al.* Exercise-induced promotion of hippocampal cell proliferation requires beta-endorphin. *FASEB J* **22**, 2253-2262, doi:10.1096/fj.07-099101 (2008).

9 Koziris, L. P. *et al.* Serum levels of total and free IGF-I and IGFBP-3 are increased and maintained in long-term training. *J Appl Physiol (1985)* **86**, 1436-1442, doi:10.1152/jappl.1999.86.4.1436 (1999).

10 Laufs, U. *et al.* Physical training increases endothelial progenitor cells, inhibits neointima formation, and enhances angiogenesis. *Circulation* **109**, 220-226, doi:10.1161/01.CIR.0000109141.48980.37 (2004).

11 Leiter, O. *et al.* Exercise-Induced Activated Platelets Increase Adult Hippocampal Precursor Proliferation and Promote Neuronal Differentiation. *Stem Cell Reports* **12**, 667-679, doi:10.1016/j.stemcr.2019.02.009 (2019).

12 Moon, H. Y. *et al.* Running-Induced Systemic Cathepsin B Secretion Is Associated with Memory Function. *Cell Metab* **24**, 332-340, doi:10.1016/j.cmet.2016.05.025 (2016).

13 Schabitz, W. R. *et al.* Intravenous brain-derived neurotrophic factor reduces infarct size and counterregulates Bax and Bcl-2 expression after temporary focal cerebral ischemia. *Stroke* **31**, 2212-2217 (2000).

14 Trejo, J. L., Carro, E. & Torres-Aleman, I. Circulating insulin-like growth factor I mediates exercise-induced increases in the number of new neurons in the adult hippocampus. *J Neurosci* **21**, 1628-1634 (2001).

15 Wrann, C. D. *et al.* Exercise induces hippocampal BDNF through a PGC-1alpha/FNDC5 pathway. *Cell Metab* **18**, 649-659, doi:10.1016/j.cmet.2013.09.008 (2013).
