## Extended data table 2 for "Exercise conditioned plasma dampens inflammation via clusterin and boosts memory"

| Patient demographics | |
|---|---|
| Age, mean (SD), years | 69.68 ± 7.95 |
| Gender male, n (%) | 19 (95) |
| number per group | 20 |
| Married, n (%) | 15 (75) |
| Education above high school, n (%) | 18 (90) |
| Emploed, full-time, n (%) | 4 (20) |
| Ethnicity, Not Hispanic, n (%) | 15 (75) |
| Race (white), n (%) | 15 (75) |
| BMI, mean (SD), kg/m2 | 27.71 (3.76) |
| Diagnose - APOE | |
| 2/3 | 3 (15) |
| 2/4 | 0 |
| 3/3 | 14 (70) |
| 3/4 | 3 (15) |

**Extended data table 2.** Patient demographics
