## Extended data table 3 for "Exercise conditioned plasma dampens inflammation via clusterin and boosts memory"

| Antibody | Species | Concentration | Manufacturer | Catalogue number | Manufacturer application recommendations and validation |
|---|---|---|---|---|---|
| BrdU | Rat | 1:2500 | Abcam | ab6326 | This antibody reacts with BrdU (5-Bromo-2'-deoxyuridine) in single stranded DNA, BrdU attached to a protein carrier or free BrdU. It detects nucleated cells that have had BrdU incorporated into their DNA during S phase. |
| GFAP | Mouse | 1:1000 | Millipore | MAB360 | Species reactivity: mouse. Tested in human brain tissue. We control tested this antibody using negative controls (no primary antibody). |
| SOX2 | Goat | 1:1000 | Santa Cruz Biotechnology | sc-365823 | Recommended for detection of Sox-2 of mouse. Control tested in human esophagus tissue and C6 and 293 cell lysates. We control tested this antibody using negative controls (no primary antibody). |
| NeuN | Mouse | 1:1000 | Millipore | MAB377 | Species reactivity: mouse. Tested in mouse dentate gyrus and rat cerebelum. |
| Doublecortin | Goat | 1:500 | Santa Cruz Biotechnology | sc-8066 | Recommended for detection of Doublecortin of mouse. We control tested this antibody using negative controls (no primary antibody) and comparison to already published doublecorting stainings. |
| Clusterin | Rabbit | N/A | Abcam | ab184100 | Species reactivity: mouse. Tested in mouse serum, lung and adrenal gland lysate. Control test in rats. |
| PEDF | Goat | N/A | R&D Systems | AF1149 | Species reactivity: mouse. Tested in C2C12 , NIH‑3T3 and 3T3-L1 mouse cell lines. |
| Factor H | Sheep | N/A | Abcam | ab8842 | Species reactivity: mouse. Tested in mouse serum. |
| LIFR | Rabbit | N/A | Proteintech | 22779-1-AP | Species specificity: mouse. Tested in mouse and human skeletal muscle tissue |

**Extended data table 3.** Primary antibody information
